# Un–complicating protein complex prediction

**DOI:** 10.1101/017376

**Authors:** Konstantinos Koutroumpas, François Képès

## Abstract

Identification of protein complexes from proteomic experiments is crucial to understand not only their function but also the principles of cellular organization. Advances in experimental techniques have enabled the construction of large–scale protein–protein interaction networks, and computational methods have been developed to analyze high–throughput data. In most cases several parameters are introduced that have to be trained before application. But how do we select the parameter values when there are no training data available? How many data do we need to properly train a method. How is the performance of a method affected when we incorrectly select the parameter values? The above questions, although important to determine the applicability of a method, are most of the time overlooked. We highlight the importance of such an analysis by investigating how limited knowledge, in the form of incomplete training data, affects the performance of parametric protein–complex prediction algorithms. Furthermore, we develop a simple non–parametric method that does not rely on the existence of training data and we compare it with the parametric alternatives. Using datasets from yeast and fly we demonstrate that parametric methods trained with limited data provide sub–optimal predictions, while our non–parametric method performs better or is on par with the parametric alternatives. Overall, our analysis questions, at least for the specific problem, whether parametric methods provide significantly better results than non–parametric ones to justify the additional effort for applying them.

## 1 Introduction

Deciphering how proteins assemble into complexes is an essential step towards better understanding their biological function. Recent improvements in high-throughput Mass Spectrometry coupled with Affinity Purification (AP–MS) technology have enabled genome–wide discovery of protein interactions and complexes in near physiological conditions [1]. In contrast to experimental methods that search for physical interactions, like yeast two–hybrid (Y2H), AP–MS directly tests complex co–membership by identifying through mass spectrometry the proteins (preys) that co–purify with a tagged protein (bait). However, several nonspecific interactors (contaminants) also co-purify with baits [1], making protein complex detection a challenging task.

During the last years, several computational methods have been developed to systematically identify complexes from such high–throughput data. Depending on the method that is employed, a different procedure has to be followed (Fig. 1). In most cases a reliability score [2–10] is initially used to discriminate between contaminants and bona fide hits. The scores assign a confidence of interaction to protein pairs based either on their co-purification similarity [2–5] or on quantitative information, *e.g.* spectral counts [6–9], MS1 intensity data [10]. Often, a binary protein interaction network (PIN) is constructed from protein interactions with scores above a pre-determined threshold, and a, usually parametric, graph-clustering algorithm is tuned and finally used to find densely connected regions (green flowchart in Fig. 1). Community/Cluster detection is a long–studied problem and a plethora of graph–clustering algorithms have been proposed. The reader is referred to [11–13] for an extended review. Recently, it has been shown that taking into account the reliability scores can improve the detection of protein complexes [14]. Nevertheless, the network has still to be filtered using a score threshold. Clearly, the value of the selected threshold significantly affects the integrity of the network and the predicted complexes. In most cases, manually curated protein-complex or protein interaction datasets are used both to choose an appropriate threshold [5], and the optimal parameters.

**Figure 1:**
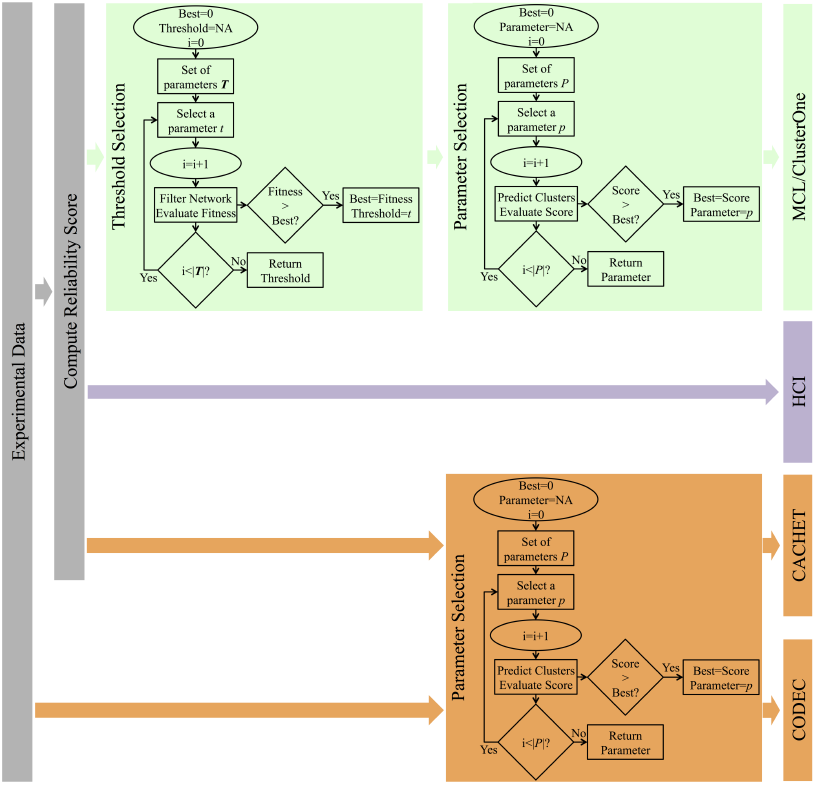
Protein complex prediction procedure for different types of computational methods. The steps that have to be taken for the prediction of protein complexes from large–scale AP-MS experiments using: (a) graph–clustering algorithms (green), (b) methods that do not construct a PIN (orange), and (c) the HCI nonparametric graph–clustering algorithm (purple).

To avoid defining a score threshold, several computational approaches [15–18] have been proposed for the prediction of protein complexes directly from AP-MS data without constructing a PIN (orange flowchart in Fig. 1). In Rungsarityotin et al., 2007 [16] a statistical model, called Markov Random Fields (MRF), is used to find the complexes that maximize the likelihood of the observations. Although several parameters are introduced, their values are directly estimated from the data, using Maximum Likelihood Estimation, and thus the method does not rely on the existence of a reference set of complexes. In the other two studies [15, 17, 18] AP–MS data are represented by bipartite graphs. A bipartite graph is a graph with two sets of vertices *U* and *V*. Each edge connects a vertex in *U* to one in *V*. In the case of AP–MS the two vertex sets correspond to the bait and prey proteins respectively, and the edges to the bait–prey interactions. COmplex DEtection from Coimmunoprecipitation data [15] (CODEC) and Core-AttaCHment structures from bipartitE TAP data [17] (CACHET), identify complexes by searching for densely connected subgraphs starting from bicliques or maximal bicliques. A biclique is a bipartite graph such that any two vertices *u* ∊ *U* and *v* ∊ *V* are connected. A biclique is maximal if it is not included in a larger biclique. Such methods have been shown to outperform classical graph–clustering methods when applied to unfiltered data. However, the superior performance of novel algorithms usually comes with an increased number of parameters that are not biologically supported and have to be optimally selected using training data or computationally expensive algorithms.

Parameter and threshold selection are challenging tasks when training data are unavailable, while improper selection may considerably affect the quality of the network and the predictions. Moreover, the computational time for the parameter optimization is proportional to the number of parameters. Assuming a search space of *M* values for each parameter the number of analyses that have to be conducted is *M*^*N*^, where *N* is the number of parameters. Finally, different parameter values are usually optimal for different experimental data, making it difficult to use the same parameter values in different studies.

In this study, we aim to investigate how incomplete training data affect the predictions of parametric protein-complex prediction algorithms, and if an easily conceivable nonparametric algorithm could provide results comparable to parametric methods without filtering the data. We evaluate the performance of two parametric graph-clustering algorithms: Markov CLuster algorithm (MCL) [19], and Clustering with Overlapping Neighborhood Expansion (ClusterONE) [14], when they are trained using different subsets of the gold-standard complexes. Their performances are compared to two methods developed specifically for the analysis of large–scale AP–MS data: Core–Attachment structures from bipartite TAP data (CACHET) [17], and Complex Detection from Coimmunoprecipitation data (CODEC) [15]. In addition, we present Hierarchical Clique Identification (HCI), a completely nonparametric graph-clustering algorithm. HCI searches for complexes based on two principles: (i) complexes should correspond to fully connected subgraphs (cliques), as their members should appear together in the experiments and (ii) the higher the score between two proteins, the higher the probability to be members of the same complex. In a first step the algorithm merges proteins in clusters in a greedy manner based on their interaction scores. Scores of intra-cluster interactions are then compared to protein-cluster interaction scores to identify proteins belonging to more than one cluster.

For the robustness analysis we use two large–scale AP–MS datasets [3, 4] from *Saccharomyces cerevisiae*. Although these datasets date back to 2006, they are ideal for the purposes of our study. *S. cerevisiae* is one of the best-studied organisms for which manually curated complex sets are available. We also evaluate the performance of the graph-clustering algorithms on a recent dataset from *Drosophila melanogaster*. This dataset is closer to modern AP-MS experiments, which employ quantitative proteomics to discriminate true interaction partners from contaminants. Unexpectedly, the non-parametric algorithm performs equally or better in comparison to the parametric ones, while its performance is not affected by the training data, which is the case for the parametric alternatives.

The remainder of this paper is organized as follows. A detailed description of the HCI algorithm is given in the “Methods” section, followed by a short description of the parametric algorithms with which it is compared. The evaluation methods that were used to assess the quality of the predicted complexes are also presented in “Methods". The results of the sensitivity and the robustness analysis are discussed in the “Results” section, which concludes with the application of the graph–clustering algorithms to the quantitative AP–MS data. The major conclusions of the study are highlighted in the “Discussion” section, accompanied by a discussion on how future studies can profit from our findings.

## 2 Materials and Methods

### 2.1 Hierarchical Clique Identification

#### Extraction of core complexes

The rationale behind HCI is similar to the one of hierarchical agglomerative clustering (HAC) methods. In agglomerative approaches, each object is initially assigned to its own cluster and pairs of clusters are iteratively merged until all objects are in the same cluster. The selection of clusters to be combined is based on a measure of distance between pairs of objects and one between pairs of clusters. In both cases distance is a measure of dissimilarity. A popular object distance is the Euclidean distance. In the case of clusters, the distance is computed as a function of the pairwise distances of objects in the clusters, the so–called linkage criterion. Some commonly used linkage criteria are the maximum, minimum or mean of all pairwise distances between pairs of objects, one from each cluster. At each iteration: (i) the pair of clusters with minimum distance is merged into a new cluster and (ii) the distances between the new cluster and the unchanged clusters are computed according to the linkage criterion. The procedure stops when there are no more clusters to merge.

It is important to note that in HAC one and only one pair of clusters is merged at each iteration, even in the case that there are several cluster pairs with the same distance. Moreover, the result of hierarchical clustering is a dendrogram, a tree diagram representing the hierarchical succession of the cluster merging. A set of clusters can be extracted by stopping the process when some criteria are fulfilled, *e.g.* maximum number of clusters or maximum acceptable distance of clusters to be merged. Finally, in clusters extracted by HAC methods there are no proteins belonging to more than one cluster (non–overlapping clusters). This is a limitation of HAC, as proteins may perform different biological functions as members of different complexes [20].

A novelty of HCI compared to HAC relies on its capability to support overlapping clusters. The idea to extend HAC to support overlapping clusters is not new. In a previous study [21] an extension of HAC called hierarchical agglomerative clustering with overlap (HACO) was proposed. While HACO predicts overlapping clusters it does so by introducing a margin parameter that enforces cluster overlap. Moreover, the final outcome of HACO is a dendrogram and a cutoff threshold has to be selected. In fact the authors use a cross–validation technique to train the two parameters. Our aim was to develop a completely nonparametric algorithm. To achieve this, we altered the way some of the main steps of the HAC algorithm are executed.

In HCI similarities between clusters are summarized in a weighted graph. Thus, we start by considering a weighted graph *G* = (*V, E, d*), where *V* is a set of nodes, *E* a set of edges, and *d* : *E* → ℝ a weight function (Fig. 2a). In this graph, nodes correspond to clusters while edge weights, in contrast with HAC, are measures of similarity. At each iteration, analogously to HAC, the algorithm selects the clusters to be combined and then updates the weights of the edges between clusters. For the selection of the clusters to be combined, an unweighted network *G*′ = (*V*, *E*′) is constructed. The network consists of the edges with a weight equal to the maximum weight:

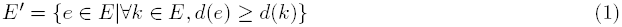

**Figure 2:**
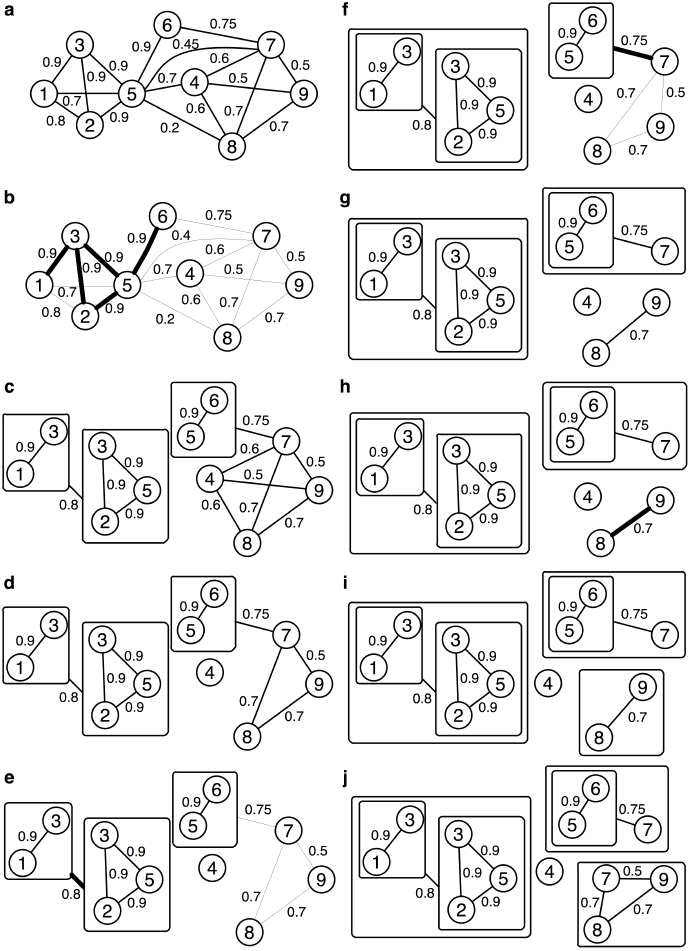
Graphical illustration of the HCI algorithm. Starting with a weighted graph (a) the algorithm creates an unweighted graph consisting of the edges with weights equal to the maximum weight (bold lines in b). The unweighted graph is mined for maximal cliques whose nodes are merged in the same cluster. Clustered nodes are removed from the network and new nodes are introduced for each cluster (c). In the case that a node loses its best neighbor (node 4 in c) then it is removed from the network (d). The new maximum weight is selected and the process continues until the network becomes disconnected (i). Nodes that may belong to additional clusters are identified and assigned as cluster attachments (j).

Then the algorithm mines this network for maximal cliques, *i.e.* cliques that are not contained in larger cliques (Fig. 2b). The extracted maximal cliques define the set of clusters to be combined. In this way, it is guaranteed that all pairs of clusters with a weight equal to the maximum weight will be merged. Moreover, given the overlapping nature of cliques, the new clusters may also overlap. Nodes corresponding to merged clusters are then removed from the network and new nodes are introduced for the new clusters (Fig. 2c). In general, linkage criteria similar to the ones used in HAC could also be used in HCI. However, the fact that clusters arisen during the execution of the algorithm may overlap should be taken into account. The linkage criterion we used was based on the two underlying principles: complexes should correspond to fully connected subgraphs, and the higher the score between two proteins the higher the probability that they are members of the same complex. Towards this end, clusters are connected if the union of their members forms a clique and the weight of the edge connecting them is equal to the maximum weight of the edges connecting their non–common members. Formally, the linkage criterion between two clusters *X* and *Y* is given by:

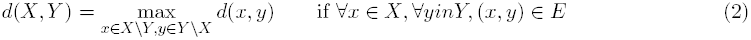

It is clear that only pairs of clusters that form a clique upon merging are connected in the new graph. This may result in the loss of the highest weighted edges of some nodes. This happens for instance when the best neighbor of a node is merged in a cluster, which is not fully connected with the node (*e.g.* node 4 in Fig. 2c). To avoid merging of the node based on lower weighted edges, which would violate our second principle, whenever a node loses its highest weighted edges it is removed from the graph (Fig. 2d). The entire procedure terminates when the graph becomes disconnected (Fig. 2i).

#### Identification of overlapping proteins

Proteins belonging to more than one complex pose a challenge to most graph clustering algorithms. Following the previously described procedure, a protein will be assigned to more than one complex only if it interacts with exactly the same score with members of different complexes. The above criterion is very strict if one considers the noise in the experimental data, the differences in the complex sizes, or the different experimental coverage of each complex (i.e fraction of complex members that have been tagged). Therefore, we assume that a protein that has already been assigned to a complex may still belong to another complex if its interactions with the members of the complex have a score comparable to the intra–cluster interaction scores. Starting with the set of complexes predicted by the above procedure, we further check whether clustered proteins could be members of additional clusters. More precisely, for a cluster *X* we define its diameter by: *r*(*X*) = min_*i∊X*_ max_*j∊X\{i}*_ *d*(*i, j*) and the similarity of a node *x* to a cluster *Y* by:

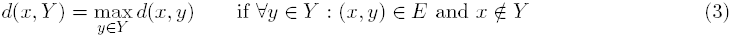

The diameter of a cluster corresponds to the weight level at which the last merging was realized in the first step of the algorithm, while similarity corresponds to the weight of the edge connecting a node to a cluster. For each cluster, nodes with similarity higher than the diameter of the cluster are identified and assigned as cluster attachments (*e.g.* node 7 in Fig. 2j). This second step of the algorithm leads to a new set of clusters with enhanced overlapping.

### 2.2 Complex prediction algorithms

**CACHET [17]** represents experimental data using weighted bipartite graphs. Initially the maximal bicliques are extracted. Bicliques that do not fulfill some criteria (minimum number of baits, minimum number of preys, minimum reliability) are discarded. A set of non–overlapping bicliques is extracted by merging bicliques with an overlap above a threshold and maximizing a reliability criterion. The member of this so–called maximum weighted independent set (MWIS) are extended with attachment proteins that are selected based on the connectivity of each protein with the cluster and the average reliability score of the connecting edges. The method introduces five parameters. CACHET has been applied on the two yeast datasets using three different scoring schemes. For both datasets, the best performance was achieved with the SAI scores. The corresponding sets of predicted complexes are available in http://www1.i2r.a--star.edu.sg/∼xlli/CACHET/CACHET.htm and they were used in this study for benchmarking.

**CODEC [15]** uses an unweighted bipartite graph to model AP–MS purifications. The method starts by mining the graph for bicliques. The bicliques are then evaluated by a likelihood score that measures the fit of the biclique to a protein complex model versus the chance that the subgraph arises at random. The highest scoring bicliques are used as seeds in a greedy expansion step that adds and deletes vertices to the biclique aiming at the identification of the best scoring biclique. The process is re–applied to random graphs with the same vertex degree as the initial bipartite graph, and a p–value is assigned to each cluster. Clusters with a p–value above a threshold are discarded. Finally, the similarity (overlap) between the remaining clusters is measured and if the similarity between two clusters exceeds a predefined threshold, the cluster with the higher pvalue is discarded. As in CACHET five parameters are introduced, however some of them can be estimated from the data. The sets of protein complexes predicted by CODEC for the yeast datasets were downloaded from http://www.cs.tau. ac.il/∼roded/CODEC/main.html.

**MCL [19]** aims at the identification of graph clusters by simulating random walks in the graph. The principle of MCL is that when a random walk reaches a densely connected cluster it will remain in the cluster until many of its vertices have been visited. MCL is simple to use, and there is only one parameter to be selected. It has been shown to be very robust and effective in identifying high–quality graph clusters [22] in comparison to other methods. The main drawback of MCL is that predicted complexes cannot overlap. In reality protein complexes often overlap and shared proteins may perform different biological functions as members of different complexes [20]. Nevertheless, MCL has been employed in many studies [8, 23–25] for the identification of protein complexes from the noisy networks constructed by AP–MS data. The single parameter of MCL was optimized by trying values from 1.8 to 5 with a step of 0.2.

**ClusterONE [14]** is based on a score, called cohesiveness, that measures how likely it is for a group of proteins to form a protein complex according to two principles: a subgraph representing a protein complex should contain many reliable interactions between its subunits, and it should be well-separated from the rest of the network. Sub–graphs with high cohesiveness are identified by an iterative greedy procedure. Finally, highly overlapping groups are merged, and groups with density below a given threshold are discarded. The method introduces three parameters: a penalty term that models the uncertainty in the data by assuming the existence of yet undiscovered interactions in the protein interaction network, an overlapping threshold, and a density threshold. In our analysis the penalty term was selected from the set {0, 1, 2, 3, 4, 5}, the overlapping threshold from the set {0.6, 0.7, 0.8, 0.9}, and the density threshold from the set {0.005, 0.01, 0.025, 0.05, 0.1, 0.2, 0.3, 0.4, 0.5}.

The methods were selected based on their performance in recent studies. Evaluation of the complexes predicted by CODEC using the yeast datasets [15] indicated that CODEC provides slightly better results than the MRF method introduced in [16], while a similar analysis for CACHET showed that CACHET outperforms CODEC [17]. ClusterOne was applied on four networks constructed from AP–MS data. It was shown that it provides the best results in comparison to seven commonly used graph–clustering algorithms, followed by MCL [14].

### 2.3 Experimental Data

Experimental data from three AP–MS studies, two [3, 4] on *S. cerevisiae* and one [8] on *D. melanogaster*, were used for the evaluation of the algorithms. In each experiment, different methods were used for protein identification by mass spectrometry. Gavin et al. [3] used matrix–assisted laser desorption/ionization–time of flight (MALDI–TOF) MS to identify proteins present in the purification. Guruharsha et al. [8] used liquid chromatography tandem MS (LC–MS/MS). Krogan et al. [4] used MALDI–TOF and LC– MS/MS, in an attempt to increase the coverage and the confidence of the observed interactome by combining the two methods. Due to the different techniques employed, the studies also differ on the level of bait coverage. Bait coverage can be defined as the fraction of all prey proteins that were also used as baits (bait–prey ratio). The first yeast dataset from Gavin et al. [3] contains 1993 bait proteins and 2671 prey proteins. The second dataset from Krogan et al. [4] contains 2294 bait proteins and 5333 prey proteins. The fly dataset from Guruharsha et al. [8] contains 3313 bait proteins and 4927 prey proteins.

#### PPI network construction

In each study a different reliability score was used to transform experimental data into weighted protein interaction networks. More specifically, Gavin et al. used socio– affinity index [3] (SAI), Krogan et al. used purification enrichment [2] (PE) and Guruharsha et al. used HyperGeometric Spectral Counts score (HGSCore) [8]. Except for the different approach each method uses to score protein interactions, there is also a striking difference in the nature of the data used. While SAI and PE use binary co–occurrence observations to score protein interactions, HGSCore exploits the quantitative aspect of the data by taking into account total spectral count, the total number of spectra identified for a protein that has been shown to correlate with protein abundance in a sample.

#### Weight normalization

All three scores have different intervals, and some of them may take negative values. Given that some algorithms do not accept negative weights, all scores were constrained between zero and one. More precisely, all scores were normalized according to:

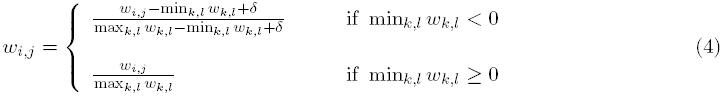

where *δ* is a small positive number (0.01) that is introduced to guarantee that interactions with minimum weight will not be assigned zero weight.

### 2.4 Clustering evaluation

#### Reference complexes

A straightforward way to evaluate predicted complexes is through direct comparison with known complexes (gold–standard). The most widely used “gold–standard” protein complex dataset for the budding yeast *S. cerevisiae* has been the manually curated protein complexes of the Munich Information center for Protein Sequences (MIPS) database [26]. However, *S. cerevisiae* protein complexes in MIPS have not been updated since 2006, lacking newly identified yeast complexes. Here, a more recent protein complex catalogue (denoted as CYC2008) was used instead of MIPS. CYC2008 [27]includes manually curated yeast complexes derived in small–scale studies and complexes identified in recent high–throughput studies that are also supported by small–scale experiments. The final list is more complete, consisting of 408 complexes in comparison to the almost 200 complexes of MIPS catalogue.

Contrary to *S. cerevisiae*, there are no hand–curated protein complex datasets for the fly *D. melanogaster*. The only fly protein complex dataset that we are aware of is one [28]that was recently compiled by integrating literature curated and computationally predicted complexes for three organisms, human, yeast and fly. Literature annotated complexes were collected from various public resources for each organism and extracted complexes were mapped between organisms using an ortholog–mapping tool. Moreover, protein complexes were predicted by applying complex prediction tools to PINs compiled by PPI databases, organism–specific databases and high–throughput datasets. From the around 6500 *D. melanogaster* complexes, less than half (≈ 3000) are literature based and the majority of them (≈ 2300) has been mapped from human and yeast. In addition, from the remaining 700 complexes, more than 500 are complexes identified in the AP–MS study [8]we also used. Given that Guruharsha et al. [8]used MCL to predict protein complexes, we decided to exclude these complexes from our reference set, as this would favor MCL results. Predicted complexes were also excluded, as they have not been experimentally confirmed in any organism. Our final reference dataset (denoted as COMPLEAT) consists of 2383 complexes. Almost 50% (1115) of these complexes are redundant, comprising complexes that either are subsets of other complexes (871) or differ from other complexes by only a few components (at least 80% of the complex is part of another complex). Only 93 complexes have no overlap with any other complex. We preserved such redundancies as suggested in the original study [28].

It is important to note that, before we evaluate predicted complexes, reference complexes were mapped to the unfiltered PIN. Known complexes with less than two proteins in the network were removed, while for the rest a new complex was defined by the subset of proteins in the complex appearing in the network.

#### Matching scores

The overlapping nature of known and predicted complex sets poses a challenge to measures that try to evaluate the matching of predicted complexes to known ones. For instance, a predicted complex may (partially) match more than one reference complex and vice versa. Several measures have been proposed for the fair evaluation of predicted complexes. However in most studies a combination of measures is used since a single measure may not be adequate. In the present study, predicted complexes are evaluated by a composite score that is computed by the geometric mean of two matching scores, the geometric accuracy [22] and the maximum matching ratio [14]. For completeness, a description of the two measures is given below.

Geometric accuracy [22] (*Acc*), is defined as the geometric mean of clustering–wise sensitivity (*Sn*) and clustering–wise positive predictive value (*PPV*). For a reference complex *i*, its complex–wise sensitivity is defined as the coverage of complex *i* by its best–matching predicted cluster: 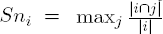, where |*i*| denotes the number of proteins in complex *i* and |*i* ∩ *j*| the number of proteins that appear both in complex *i* and predicted complex *j*. The general sensitivity of a set of predicted clusters is given by the clustering–wise sensitivity, the weighted average of complex–wise sensitivity over all complexes: 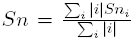. Similarly to complex–wise sensitivity, the complex–wise positive predictive value represents the maximal fraction of proteins in cluster *j* found in the same reference complex: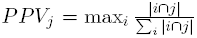 and the general positive predictive value of a set of predicted clusters is given by the weighted average of complex–wise positive predictive value over all clusters: 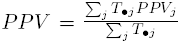, where 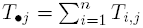. Given that *Sn* is maximized when all proteins are grouped in a single cluster, while *PPV* is maximized when each protein is assigned on its own cluster the two measures can be integrated by taking their geometric mean, which corresponds to geometric accuracy: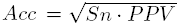. A caveat of geometric accuracy is that the denominator *T*_*•j*_ may be larger, smaller or equal to the number of proteins in the best–matching reference complex and it has be shown [14, 22] that this results in lower scores when the predicted complexes overlap. For instance, when is used to compare CYC2008, in which around 200 proteins belong to more than one complex, with itself we take a score of 0.827 instead of 1, while it gets even lower, 0.292, when the highly overlapping fly gold–standard is compared with itself.

Maximum matching ratio [14] (*M M R*), tries to overcome this problem by searching a maximum one–to–one mapping between predicted and reference complexes. A bipartite graph is formed, in which the two sets of nodes represent the reference and predicted complexes. Edges connect reference complexes with the predicted complexes with which they overlap and their weights correspond to the overlap [29]between the two complexes, given by: 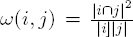. The bipartite graph is mined for a subset of edges such that each predicted and reference complex appears only in one of the selected edges and the sum of the weights of the selected edges is maximal. *M M R* is then given by the sum of the weights of the maximum weighted bipartite matching divided by the number of reference complexes. An advantage of *M M R* is that the maximum weighted bipartite matching represents an optimal assignment between reference and predicted complexes as no reference complex is assigned to more than one predicted complex and vice versa.

## 3 Results

### 3.1 Sensitivity Analysis

Initially a parameter sensitivity analysis of the two parametric graph–clustering algorithms was performed. For each *S. cerevisiae* dataset the two algorithms were applied to two PPI networks. The first network, termed “Unthresholded", is constructed by simply applying the corresponding reliability score to the experimental data, *i.e.* SAI for Gavin et al. and PE for Krogan et al. The second “Thresholded” network results after the removal of protein interactions with scores below the thresholds that were proposed in the original studies, 5 for SAI and 3.19 for PE. The results are shown in Fig. 3. For ClusterONE we found that density threshold is the parameter that mostly affects algorithms predictions. For this reason the variations in performance due to variations in the other two parameters are shown as error bars (Fig. 3b, d). It is clear that network filtering improves the performance of both algorithms. In the case of MCL the difference between the best scores attained using the thresholded and unthresholded networks is more prominent in Krogan et al. data. A possible explanation for this may be the difference in the bait–prey coverage between the two yeast datasets. On the other side, ClusterONE performance seems to be less affected by the lower bait–prey ratio. Filtering also affects the range of the performances. The variation in the scores is smaller for the thresholded networks in comparison to the unthresholded ones, and the performance of both algorithms seems to be more stable when a score threshold is applied. Finally, as we already mentioned, different parameter values are optimal for different PPI networks. Although this does not considerably affect MCL algorithm, this is not the case for ClusterONE. For instance, the optimal density threshold for the thresholded network from Gavin et al. (0.2) provides significantly lower scores not only for the two networks from Krogan et al. but also for the unthresholded network from Gavin et al. The above results clearly indicate that although the additional degrees of freedom of ClusterONE make the algorithm more flexible, *i.e.* it can be trained to provide very good results for different datasets, at the same time parameter fitting becomes more difficult.

**Figure 3:**
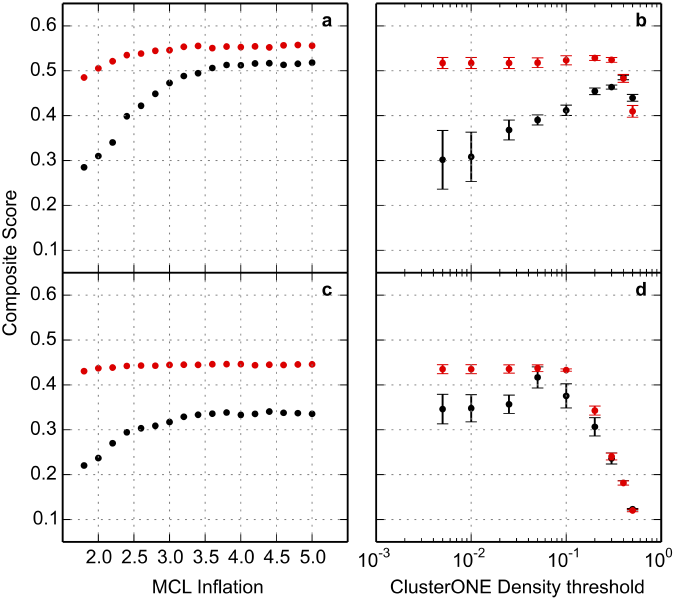
MCL and ClusterONE sensitivity analysis. Matching scores obtained for parameter variations of MCL (a, c) and ClusterONE (b, d). For ClusterONE density threshold is the parameter that mostly affects algorithms predictions. The variations in performance due to variations in the other two parameters are shown as error bars. The algorithms were applied to PPI networks constructed from the Gavin et al. (a, b) and Krogan et al. (c, d) datasets, with and without using a score threshold, red and black curves respectively.

### 3.2 Robustness Analysis

To check how limited training data affect the performance of parametric methods, random subsets of the reference complexes set were used for training. The fraction of the gold-standard complexes used for training was varied in the set {0.02, 0.03, 0.04, 0.05, 0.06, 0.07, 0.08, 0.09, 0.1, 0.2, 0.4, 0.6}. In the case of the Gavin et al. dataset the fractions correspond to subsets of {5, 8, 10, 13, 15, 18, 20, 23, 25, 50, 100, 149} complexes, while in the case of Krogan et al. dataset to subsets of {8, 12, 16, 20, 24, 28, 32, 36, 39, 78, 156, 234} complexes. Given that training subsets of the same size but consisting of different reference complexes may result in different optimal parameters and consequently in different performances, for each fraction 100 subsets were randomly selected. For a fraction *f* and a random subset *i* the parameters 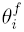 that provide the best match to the subset were found and the matching score 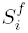 using the whole set of reference complexes was computed. In this way for each fraction *f* a set 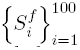 of scores is collected that can be used to compute several statistics like the expected value and the variance. Box plots of the sets of scores for the two yeast datasets are shown in Figs. 4–5. For comparison the performances of HCI, CODEC and CACHET have been overlaid.

**Figure 4:**
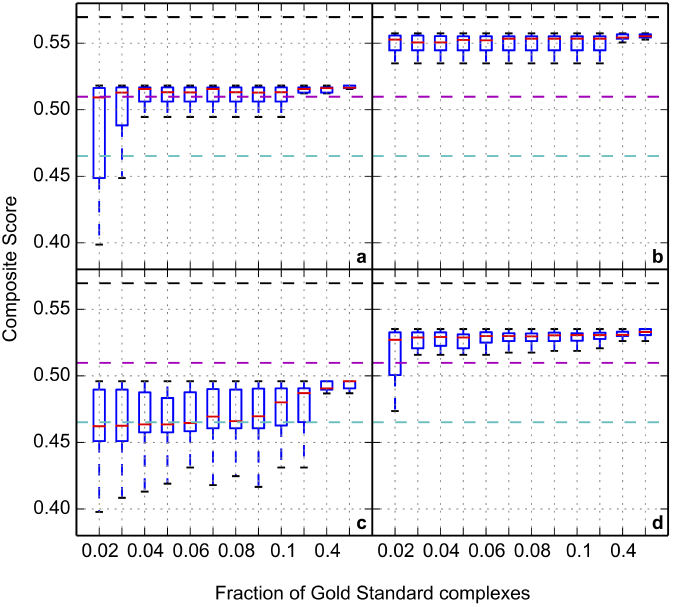
Robustness analysis using Gavin et al. dataset. Evaluation of the complexes predicted by HCI, ClusterONE, MCL, CODEC and CACHET for the Gavin et al. [3] dataset. MCL (a, b) and ClusterONE (c, d) parameters were trained using fractions of the CYC2008 complexes, and they were applied to unthresholded (a, c) and thresholded (b, d) networks. For HCI (black dashed line), CODEC (magenta dashed line), and CACHET (cyan dashed line) no threshold was used.

**Figure 5:**
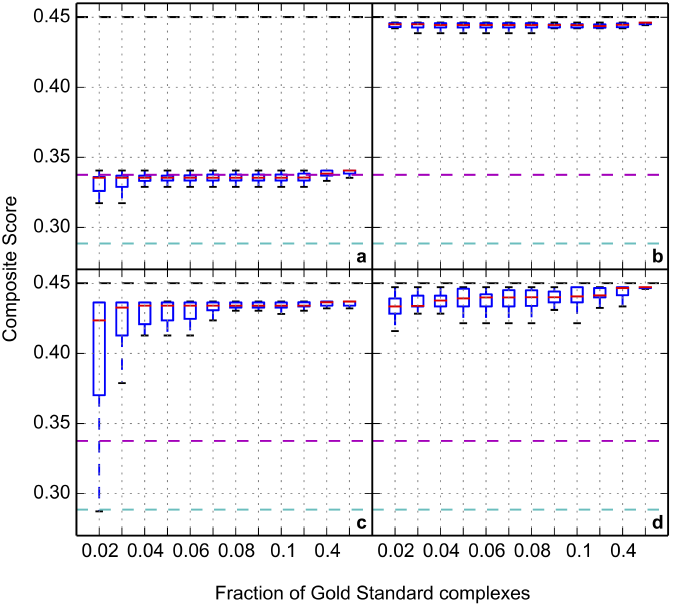
Robustness analysis using Krogan et al. dataset. Evaluation of the complexes predicted by HCI, ClusterONE, MCL, CODEC and CACHET for the Krogan et al. [4] dataset. See Fig. 4 caption.

As expected, training with limited data affects the performance of both algorithms, increasing the variability of the attained scores and decreasing the median score. The median score asymptotically converges to the optimal score as the size of the training set increases. MCL seems to be less sensitive to incomplete training data, as its median scores are closer and converge faster to the optimal score when compared to ClusterONE. This may happen either because the optimal parameter can be identified with fewer data or because there is no big difference between the scores of the selected versus optimal parameters. We also observe that the performance of both methods is less affected in the thresholded networks. This may be in part due to the more stable performance of both algorithms in thresholded networks.

Comparing with the other methods we observe that the nonparametric algorithm provides very good results. HCI gives the best predictions, and its performance is better or very close to the performance attained when the other graph–clustering algorithms are applied either on the unthresholded or on the thresholded networks. Therefore, HCI, despite being nonparametric, can provide top scoring results without the need to define a score threshold. HCI also outperforms CODEC and CACHET, which could not match the performance of graph–clustering algorithms in all cases. It is important to note that CODEC and CACHET are parametric methods, and their performance may also be affected by limited training data. Due to difficulties in their implementation and application they were not included in the robustness and sensitivity analysis.

### 3.3 Quantitative AP–MS data

The two yeast datasets have been extensively used as benchmarks in many similar studies. However, they differ from modern AP–MS experiments, which employ quantitative proteomics to discriminate true interaction partners from contaminants. We investigated how the three graph–clustering algorithms, MCL, ClusterONE, and HCI, perform on such kind of data using a more recent dataset from *D. melanogaster* [8]. Similarly to the yeast datasets, HCI predictions give the best matching scores when compared to the predictions of MCL and ClusterONE when the latter are applied on the unthresholded network (Fig. 6). The performances of both graph–clustering algorithms improve with the filtering of the data, but they remain slightly worse than HCI performance. In general, HCI predictions using unthresholded data are better or comparable to the best predictions of its competitors. The performance of all methods is considerably lower in fly data in comparison to the yeast ones. The reason is that COMPLEAT is not manually curated and consists of highly overlapping complexes that affect the measures we have selected. While highly overlapping complexes should probably be merged, they were preserved as suggested in the original study [28].

**Figure 6:**
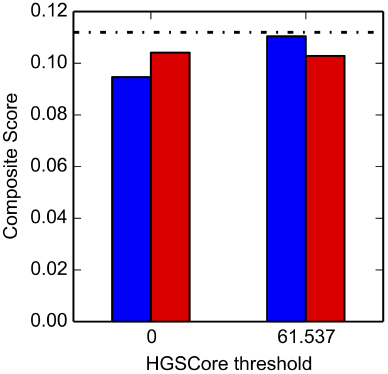
Performance on quantitative AP–MS data. Evaluation of the complexes predicted by HCI (black dashed line), ClusterONE (red bars), and MCL (blue bars) for the *D. melanogaster* [8] dataset. ClusterONE and MCL were applied to the unthresholded, tantamount to a 0 threshold, and the thresholded network.

## 4 Discussion

Computational methods play an important role in the analysis of the vast amount of data produced by recent high–throughput experiments. In many cases their application relies on the existence of biological data that are necessary for the training of a statistical or machine–learning approach or the identification of the values of the parameters that are introduced by the method. Although training data are sometimes available, this may not be the case for experimental studies focusing on unexplored biological areas/systems, for which there is limited or no previous knowledge. Using high–throughput protein interaction data and state–of–the–art protein complex prediction methods, we showed that training with limited data affects the performance of parametric approaches. One could claim that there is no reason to re–train the methods, and previously identified optimal parameters can be used in the future. However, our analysis indicates that transfer of optimal parameter values from one study to another does not guarantee optimal results, especially when the algorithm is very sensitive to the parameter value.

A way to overcome the limitations of parametric methods is to use nonparametric ones. We developed HCI, a nonparametric graph–clustering algorithm, and we compared its performance to its parametric alternatives. The results indicate that HCI can efficiently predict protein complexes from high–throughput protein interaction data without introducing unnecessary parameters, providing very good results in comparison to the other methods. Therefore, a simple nonparametric method could match the performance of the more sophisticated parametric methods.

Despite our findings, the current study should not be considered as an anathema to parametric methods. For several biological problems, e.g. the prediction of protein–complexes from small to intermediate scale AP–MS data [30, 31], the use of parameters is unavoidable, as nonparametric methods cannot provide adequate results due to the noise and the complexity of the data. Even in such cases the limitations of the parametric methods should not be overlooked. While in most studies introducing a novel computational method, its performance is compared to that of existing methods; it is seldom the case that a parameter selection policy is described, or a sensitivity or robustness analysis is conducted. An analysis similar to this study provides helpful insights on the behavior of a computational method under non–optimal conditions. We also consider important the use of simple nonparametric methods as benchmarks, as in this way the added value of a sophisticated method can be properly evaluated.

Regarding the problem of protein complex prediction from large–scale AP–MS data, HCI seems to be a good alternative to existing methods; especially in cases where parameter selection and data filtering are cumbered by the lack of training data or the limited expertise of the user. The algorithm provided very good results for three AP–MS datasets, using various confidence scores. It is important to note that the scores used in this study fall in the class of matrix approaches, which compute confidence scores both for bait–prey, and prey–prey interactions. While the method can also be used with approaches that score bait–prey interactions only, in such cases the existence of unobserved interactions in the inferred PIN should be taken into account in the interpretation of the predictions. The HCI algorithm is an extension of hierarchical agglomerative clustering that addresses some of its limitations. Given the broad applicability of hierarchical agglomerative clustering, we believe that HCI can also be applied to problems for which this family of algorithms is employed.

